# Synthesis and characterization of oral drug delivery, a pH – sensitive silver nanocomposite based on sodium alginate extracted from *Sargassum asperifolium* collected from Jazan coasts, KSA

**DOI:** 10.1101/557231

**Authors:** Fatimah. A. Agili, Sahera. F. Mohamed

## Abstract

The pH-sensitive nanocomposite composed of sodium alginate/ Pectin/ Tannic acid – silver SA/Pec/TA-Ag was prepared using microwave irradiation and employed as a carrier for Propranolol drug. Physico-chemical characteristics of the prepared systems using Fourier Transform Infrared Spectroscopy (FTIR), X-ray Diffraction (XRD), Field Emission Scanning Electron Microscope (FESEM), High-Resolution Transmission Electron Microscope (HRTEM), Dynamic light Scattering instrument (DLS), and Energy Dispersive X-Ray Analysis (EDX). The percentage drug release was 96% at pH 7.4 within 420 min. The drug release data was fitted into different kinetic models included zero order, First order, Higuchi and Ritger-Peppas model. The release mechanism is non-Fickian character where it controlled by diffusion and relaxation of polymer chains. It can be concluded that SA/Pec/TA-Ag nanocomposite is candidate for the oral drug carrier specific for intestinal system and has stability against gastric fluid.

## Introduction

Smart hydrogels are hydrophilic polymeric three-dimensional systems that have physically or synthetically cross-inked polymer. They quickly swell and contract because of natural changes, for example, pH, temperature, magnetic or electric field because of the presence of certain functional groups along the polymeric chain [1]. Imaginative medication conveyance advances in view of insightful hydrogels have extraordinary benefit [2, 3]. The pH-responsive hydrogels are considered as an appealing for the controlled release of medications because of their pH responsively moves in various locales of the body in both typical and pathological conditions [4].

Biopolymers hydrogels provide great applications in pharmaceutical field due to non-poisonous quality, high biocompatibility and biodegradability [5]. Sodium alginate (SA) is a characteristic polysaccharide of a purified carbohydrate extracted from brown seaweeds by utilizing dilute alkali [6]. It is a polyanionic natural linear copolymers of (1→4) α-Lguluronic acid (G) and (1→4) β-D-mannuronic acid (M) units it has already been widely used in a variety of biomedical applications [7]. Pectin (Pec) is natural polysaccharide found in berries, apples and other fruit. It is linear chains of α-(1–4)-linked D-galacturonic acid [8]. Tannic acid (TA) is plant polyphenol contains five pyrogallol and five catechol groups that provide multiple bonding sites with diverse interactions [9]. For this reason, it is applied on tissue engineering as a crosslinking reagent [10].

The microwave irradiation is considered a beneficial methodology for hydrogel synthesis by utilizing a mix of polymeric reactants [11]. It has high temperatures heating process for attack the solution in a short time and restricted the side reactions [12].

Nanoparticles (NP) has been extensively studied particularly noble metals because of their novel functions and exceptional properties [13]. Silver NPs are non-harmful and environmentally eco-friendly [14]. Among different noble metal NPs, silver shows extraordinary consideration due to their prevalent antibacterial properties [15]. They have high partiality toward sulfur or phosphorus containing proteins inside or outside bacterial cell layers actuating auxiliary changes, which influences bacterial cell feasibility [16, 17].

Herein, the investigation of different properties of inorganic nanomaterials incorporated biopolymer hydrogels are not uncommon, however the readiness of sodium alginate/pectin/tannic acid biopolymer hydrogel consolidated with silver nanoparticle as a drug carrier is new in the literature. In this study, a pH-responsive hydrogel of SA/Pec/TA-Ag nanocomposite was synthesized using microwave irradiation to act as a drug carrier Tannic acid here has a dual impact it acts as a crosslinker and as a reducing agent for Ag ions more over it rich in carboxylic acid groups. The nanocomposite was characterized by FTIR, XRD, FESEM, HRTEM and DLS and its pH-responsive swelling and controlled drug releasing properties were investigated.

## Materials and Methods

### Materials

Pectin (from apples) (Pec) was acquired from (Sigma-Aldrich, USA). Tannic acid (TA) supplied from (Qualikems, India). Propranolol drug was obtained from El Qahera for Pharmaceutical & Chemical Industries, Egypt. Other chemicals, for example, ammonium persulfate (APS), silver nitrate, phosphate and acetate buffers were purchased from ((Sigma-Aldrich, USA) and utilized without assist refinement.

### Algae collection

Brown alga *Sargassum asperifolium*, was in family Sargassaceae, genus *Sargassum*. It was collected from Jazan coasts, KSA. The alga was washed in distilled water, dried over night at 40–45°C in an oven. The dry weights were gained after drying overnight at 105°C.

## Methods

### Extraction and Purification of sodium alginate

The samples (10 g) were suspended in 2% CaCl2 for 2 h, washed with deionized water [18]. Alginate was extracted by addition of an aqueous solution of Na_2_CO_3_ 1 M and 0.5 g of EDTA and the pH of the suspension adjusted to pH 11 for 48 h. This was then filtered through muslin cloth. Sodium alginate was purified according to the method of Gomez *et al*., [19]. Aqueous solution of sodium alginate was directly precipitated, under stirring, by addition of ethanol until reaching a proportion 1:1 in volume, respectively. Thus, the insoluble polymer was separated and then exhaustively washed with acetone by sox let for 100 h. Finally, the biopolymer was dried at room temperature under vacuum until constant mass.

### Synthesis of SA/Pec/TA hydrogel

A constant ratios of SA: Pec: TA (10: 4: 3) were weighted according to the feed ratio and dissolved in hot water with stirring. After cooling 0.5wt% of APS (initiator) was added. The solution was briefly mixed by a prop sonication for 10 min to form a homogeneous solution. Then, it was subjected to microwave irradiation of power output 300 W for 10 min using domestic microwave oven, Thai Samsung Electronics Company. The formed hydrogel was extricated for 2h in hot water to evacuate the homopolymer and dried for 12h at 40°C in an oven.

### Synthesis of SA/Pec/TA-Ag nanocomposite

A 100 mg of dried SA/Pec/TA hydrogel was placed in 50 mL of Ag ions solution of concentration 250 mg/L for 24 h to dope metal ions in the blend matrix. Blended matrix with loaded metal ions was placed into distilled water for 24 h to remove unbound metal ions. Then, they were reduced by transferring them into 50 ml of 5% NaOH for 6 h and then in 50 ml of 0.5 M NaBH4 for another 6h to complete reduction of the metal ions. Soaking them in de-ionized water for 12 h and drying in oven at 40°C.

### Instrumentations

FTIR spectroscopy analysis was finished utilizing Nicolet Is-10 FTIR, USA, in the range of 400–4000 cm^−1^. High-resolution transmission electron microscope (HRTEM) estimations were done utilizing, JEOL 2100-Lab6, Japan. Its magnification 2000 to 1500000x. The accelerating voltage is 80 to 200 kV. Morphological examinations were done utilizing (Field Emission) SEM, Quanta 250 FEG, the accelerating voltage is 30 K.V. FEI Company, Netherlands. It attached with EDX Unit (Energy Dispersive X-beam Analyses). The XRD was done using XD-DI Series, Shimadzu apparatus with a copper target (λ = 1.542 Å). All the diffraction patterns were accomplished at operating voltage of 40 kV, an electric current of 30 mA, and at room temperature over a range of 2θ (10° – 90°) at a scan speed of 8°/min. Dynamic light scattering instrument (DLS) (Zeta Sizer Nano Series (HT), Nano ZS, Malvern Instruments, UK. Absorbance estimation was done utilizing UV/VIS spectrometer; UV-Analytic Jena AG, Germany, with a quartz cell of 1.0-cm optical length.

### The swelling measurements

The clean, dried, weighed sample was soaked in bi-distilled water or buffer solution at room temperature for different time intervals. The sample was removed and the excess water on the surface was removed by blotting quickly with filter paper and reweighed. The swelling percent was calculated as follows:

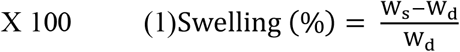

Where W_d_ and W_s_ are the masses of dry and swelled sample, respectively.

### Drug Loading

The stacking of Propranolol drug onto SA/Pec/TA hydrogel and SA/Pec/TA-Ag nanocomposite was completed by swelling harmony technique. The samples were allowed to swell in the drug solution of different concentration and pHs for 24h and afterward dried at room temperature. The concentration of rejected solution was investigated to ascertain the percent of drug caught in the lattice at the wavelength of 294nm.

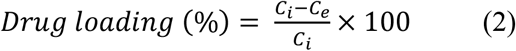

Where C_i_ (mg/L) is the initial drug concentration and C_e_ (mg/L) is the equilibrium drug concentrations in the solution.

### In vitro Propranolol drug release studies

In vitro release studies of the drug were carried out by suspending of 100 mg of the drug loaded sample in 10 mL of the buffer-releasing medium (pH 2.1 and 7.4) at 37°C. The amount of drug released was assayed by using spectrophotometer. All the studies were carried out in triplicate and the total uncertainly range was 2–4%.

### Drug release kinetic

To contemplate the Propranolol release mechanism from SA/Pec/TA hydrogel, and SA/Pec/TA-Ag nanocomposite, different kinetic models were considered to fit the experimental data. [20, 21].

Zero-order drug release kinetic model:

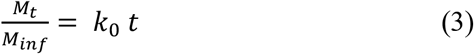

The First order drug release kinetic model:

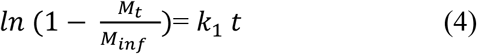

The Higuchi Square root model:

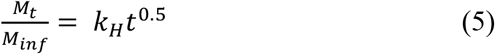

Ritger-Peppas model:

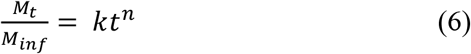

Where M_t_ and M_inf_ are the drug release at time t and equilibrium, respectively. k_0_ is zero-order release constant. k_1_ is first-order release constant. k_H_ is Higuchi release constant. k is the rate constant. n is the release exponent,

## Results and Discussion

### Physico-chemical characteristics of SA/Pec/TA and SA/Pec/TA-Ag

The FTIR spectra of SA, SA/Pec/TA hydrogel, and SA/Pec/TA-Ag nanocomposite are appeared in Fig. 1. Clearly SA demonstrates an expansive band at 3456 cm^−1^ ascribed to O-H groups. The band at 2925 cm^−1^ identified with C-H stretching vibration. The C=O stretching vibration showed up at 1543 cm^−1^. SA has a characteristic band showed up at 982 cm^−1^ because of Na–O [22]. For SA/Pec/TA hydrogel the bands of O-H, C-H, and C=O stretching vibrations showed up at 3376, 2973, and 1635 cm^−1^. The increase the intensity of O-H band and the sharpness of C=O band compared with SA because of Pec and TA present in the hydrogel network. Smaller bands were seen at 915 and 587 cm^−1^ related to the substituted benzene ring in TA [23]. For SA/Pec/TA-Ag nanocomposite, it could be noticed that no new bands were seen however; slight shifts of the bands were observed. This is because the electrostatic attraction between the Ag and the electron donating groups present in SA, Pec and TA.

**Fig. 1.**
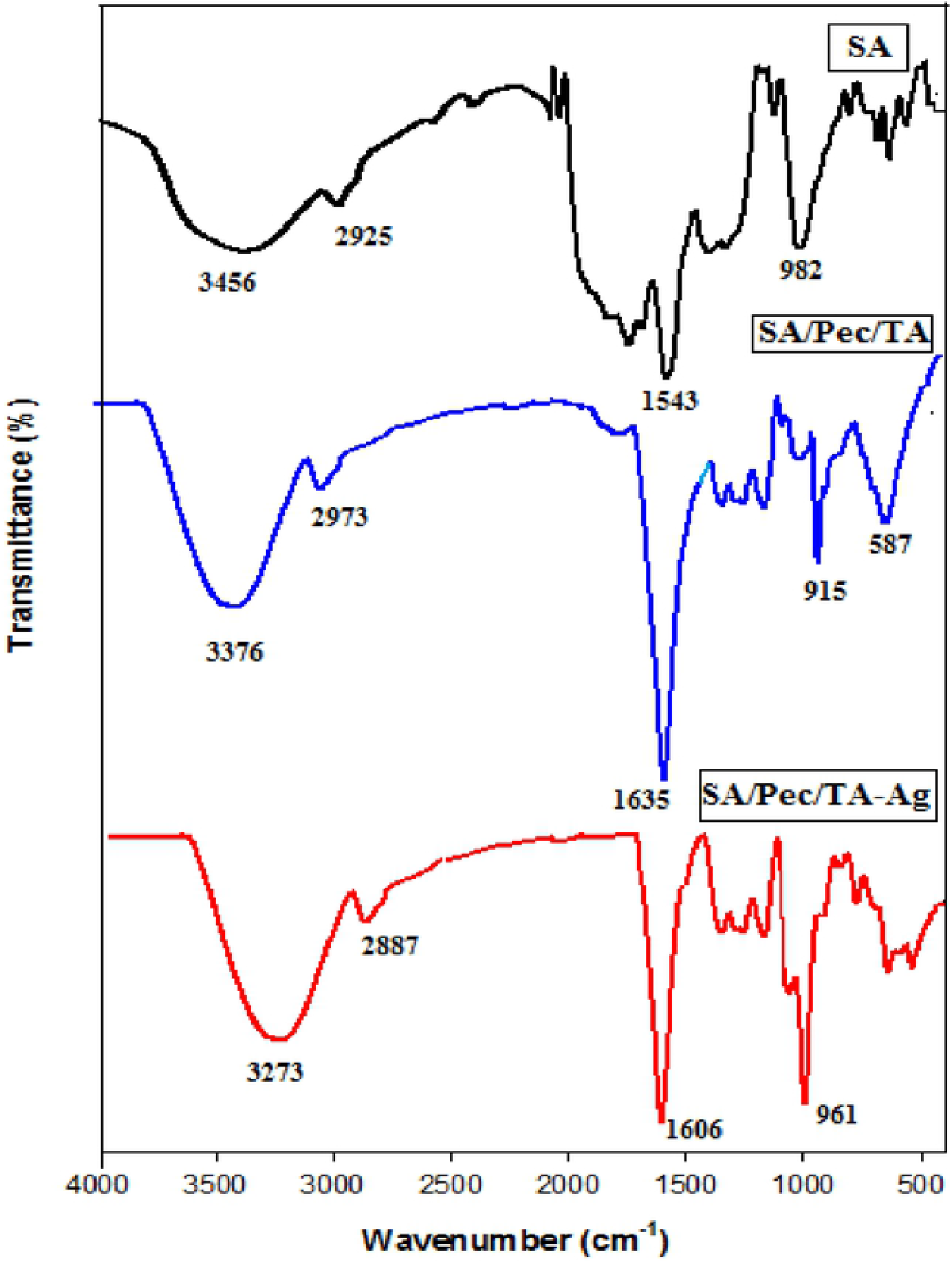
FTIR spectra of SA, SA/Pec/TA hydrogel, and SA/Pec/TA-Ag nanocomposite.

The surface morphology of SA/Pec/TA hydrogel, and SA/Pec/TA-Ag nanocomposite were inspected utilizing FESEM as appeared in Fig. 2. For SA/Pec/TA hydrogel, Fig. 2A, the surface shows up as a smooth which demonstrated good compatibility and effectively fused of the polymeric segments in the system structure of the hydrogel. For SA/Pec/TA-Ag nanocomposite, Fig. 2 B, the surface totally changed to a gruff surface with some sporadic pores on account of joining of Ag nanoparticles inside the polymeric structure. The EDX micrographs of SA/Pec/TA hydrogel, and SA/Pec/TA-Ag nanocomposite are shown in Figure 2 A′ and B′, respectively. It can be observed that C, O are the main backbone elements in the hydrogel network, Figure 2A′. The distribution of C, O, and Na in the weight percent is 40.53, 49.32 and 6.35 respectively. The presence of traces 3.8 wt% of Ca and Cl elements pointed to an elemental residue amid SA extract. For SA/Pec/TA-Ag nanocomposite, the percentage of weight of C, O, Na, and Ag is 18.02, 21.60, 2.73, and 57.65, respectively. The presence of Ag signals exhibited the presence of Ag nanoparticles.

**Fig. 2.**
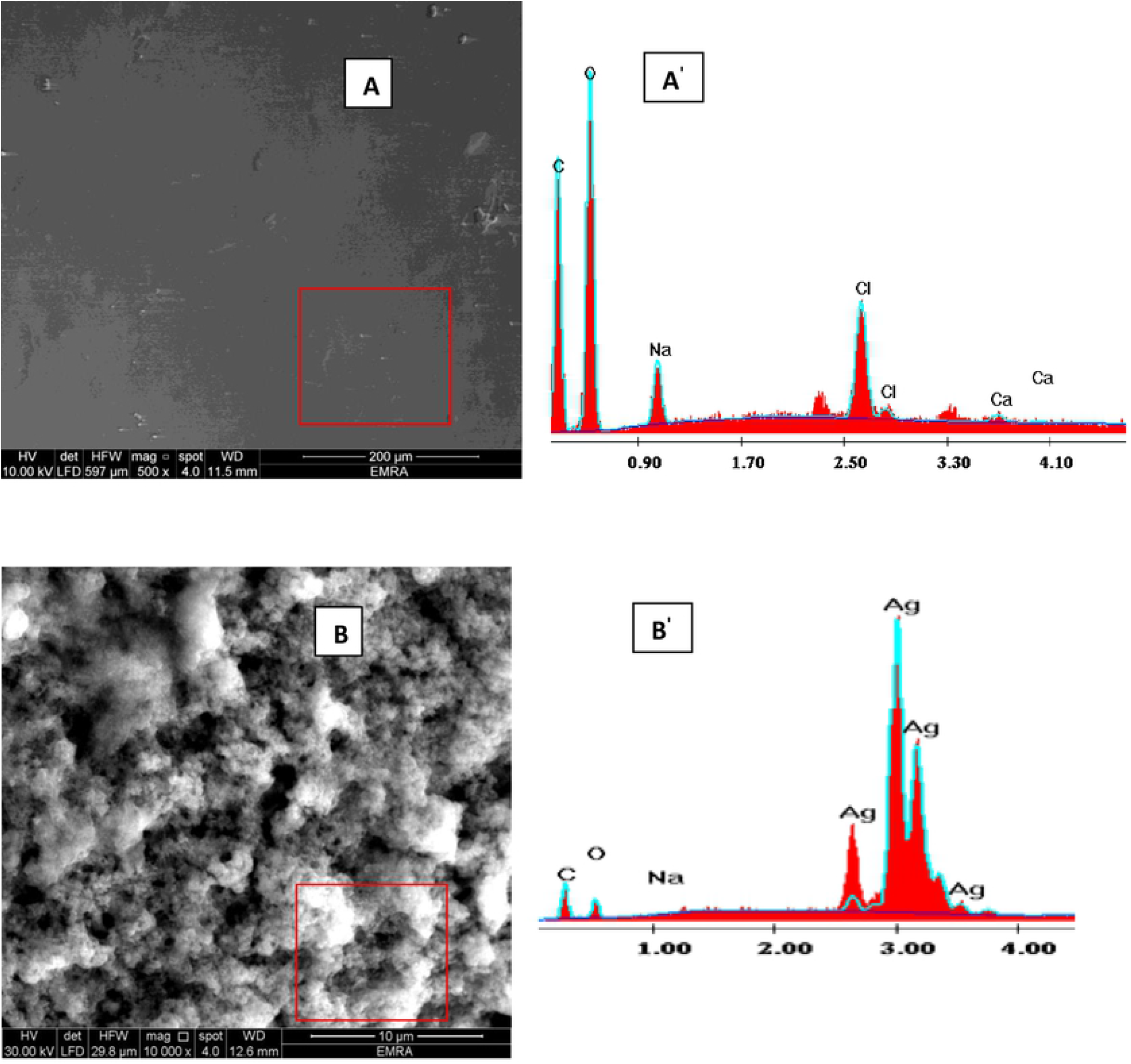
(A) FE-SEM (A′) EDX of SA/Pec/TA hydrogel, (B) FE-SEM, (B′) EDX of SA/Pec/TA-Ag nanocomposite.

Fig. 3 demonstrates the HR-TEM images of SA/Pec/TA-Ag nanocomposite. As found in the Figure, an arbitrary dispersion of Ag nanoparticles which showed up as an about circular dark of various particle size. DLS measure examination for Sa/Pec/Ag nanocomposite uncovered that the Ag NP readiness had distribution ordinarily with a noteworthy peaks of average size between 21.91 – 34.04 nm.

**Fig. 3.**
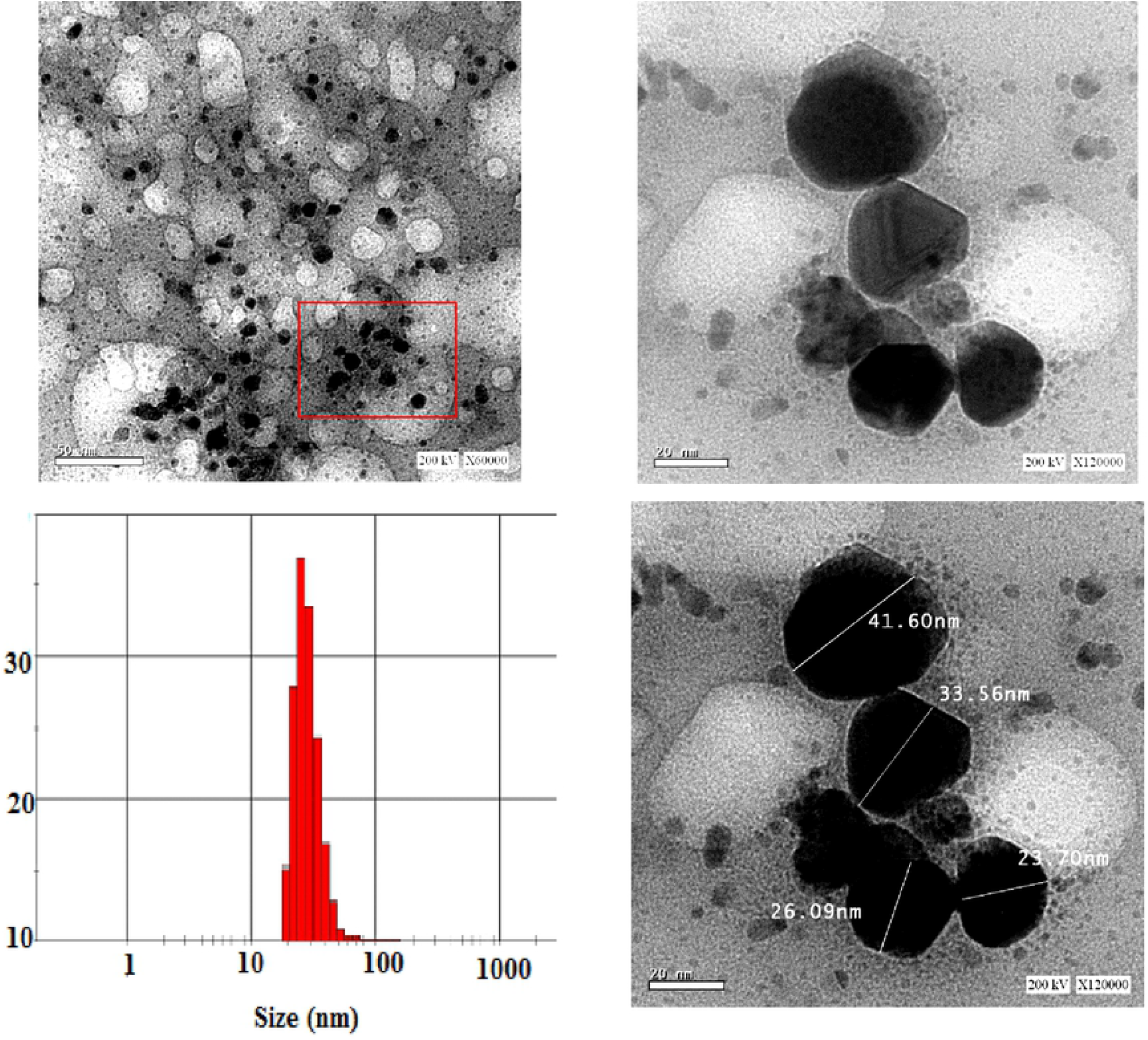
HR-TEM images of different magnification of SA/Pec/TA-Ag nanocomposite and DLS.

XRD diffractograms of SA/Pec/TA hydrogel, and SA/Pec/TA-Ag nanocomposite are appeared in Fig.4. The diffractogram of SA/Pec/TA hydrogel demonstrated a wide peak at 2θ of 19.6° demonstrating the amorphous phase of the hydrogel. The diffractogram of SA/Pec/TA-Ag nanocomposite indicates sharp diffraction peaks happened at 2θ of 37.03°, 44.46°, 64.62°, and 77.54° corresponding to (111), (200), (220), and (311) planes of the cubic Ag, respectively [24]. The presence of these peaks confirmed the presence of Ag in the nanocomposite.

**Fig. 4.**
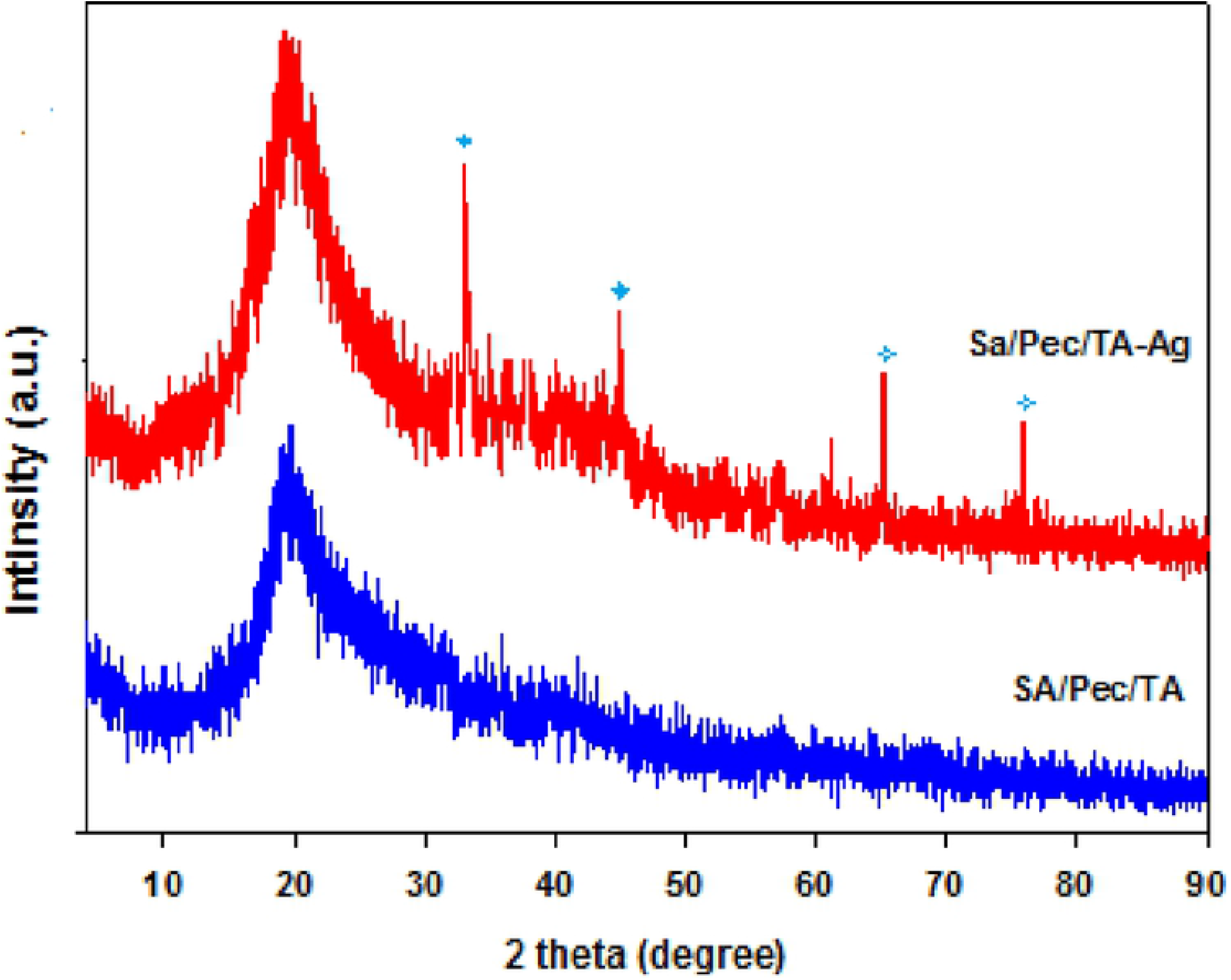
XRD diffractograms of SA/Pec/TA hydrogel, and SA/Pec/TA-Ag nanocomposite.

### Swelling behavior

The swelling behavior of SA/Pec/TA hydrogel, and SA/Pec/TA-Ag nanocomposite at various pH values as a function of time are shown in Fig 5. It can be observed that SA/Pec/TA hydrogel, and SA/Pec/TA-Ag nanocomposite exhibited higher swelling response which increased with time until equilibrium attained within 500 min. The higher swelling response attributed to the high hydrophilicity of polymeric chains. The–COOH groups in SA, Pec, and TA play a major role in swelling. When carboxylate bunches are protonated, the particles have a tendency to assimilate water to fill the pores in the polymer network, until the point that balance is accomplished.

**Fig. 5.**
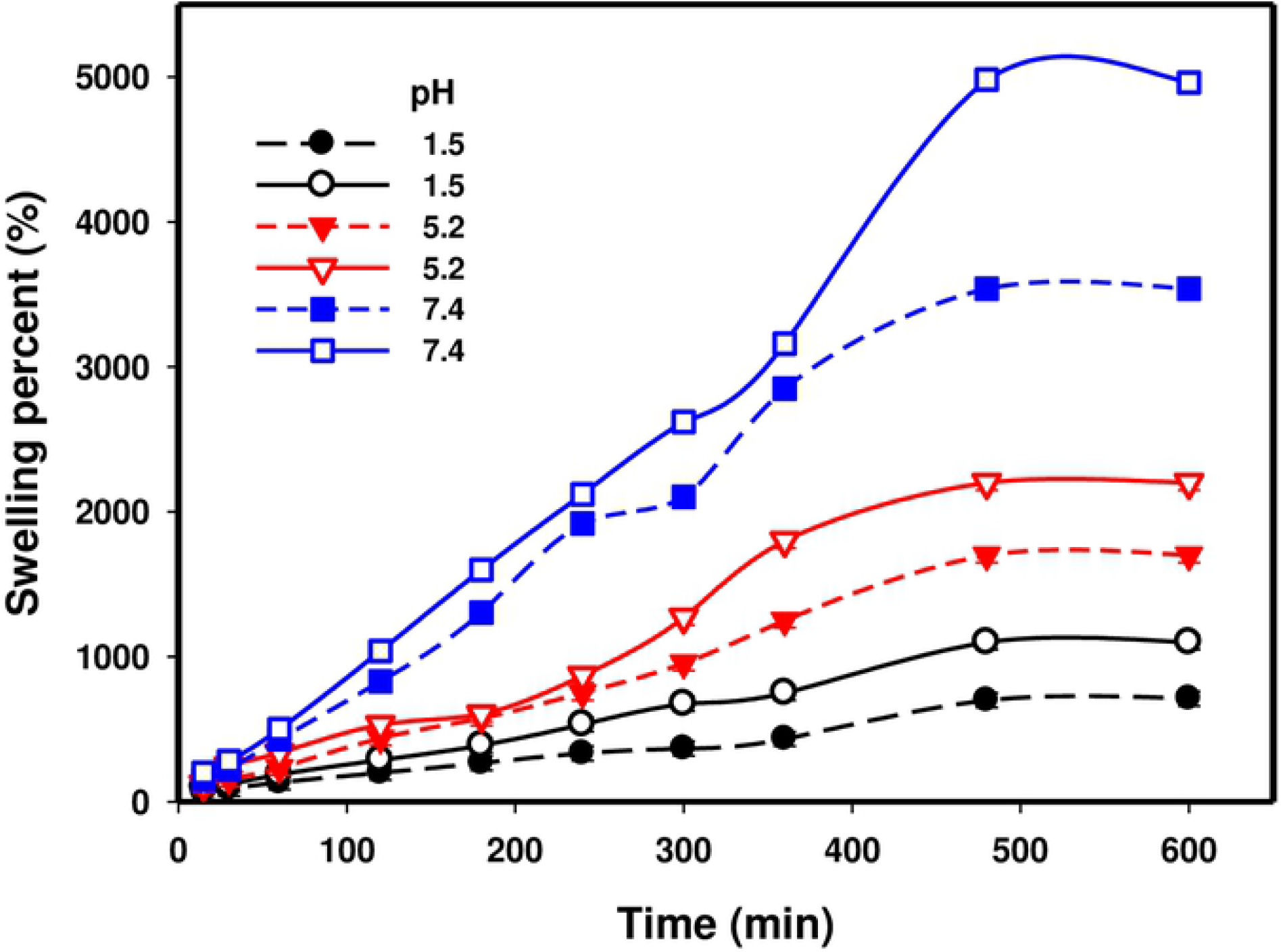
The swelling percent of SA/Pec/TA (dark symbols) hydrogel, and SA/Pec/TA-Ag nanocomposite (open symbols) as a function of time at pHs 1.5, 5.2, and 7.4.

The pH exceedingly influenced the swelling percent as obtained from Fig. 5. The swelling capacity expanded with expanding pH where the lowest pH obtained at 1.5 and the highest one obtained at pH 7.4. At pH 1.5 the carboxylic group is non-ionized. Strong hydrogen bonds formed between the polymer chains, which diminished the free volume in the hydrogel network as well as the swelling percent [25]. At pH 7.4 the −COOH groups are ionized to −COO^−^ results in a repulsion of the negatively charged on the caboxylate ions which prompts improving the free spacing in the hydrogel whatever enhancing the swelling capacity. It tends to be additionally seen that the swelling percent of SA/Pec/TA-Ag nanocomposite is higher than Sa/Pec/TA hydrogel. The nearness of Ag nanoparticles bring down crosslinking density in network structure and all the more free spaces that manage the opportunity for the rapid diffusion of water molecules into the matrix as well as enhanced the swelling behavior.

### Propranolol drug Loading

Impact of Propranolol initial concentration on the drug stacking of SA/Pec/TA hydrogel, and SA/Pec/TA-Ag nanocomposite was learned at various pHs (1.5, 5.2, and 7.4) and results are appeared in Fig. 6. It tends to be noticed that the drug loaded amount expanded with expanding the underlying drug focus because of the free volume accessible destinations on the framework allow the diffusion of more drug molecules. It must be noticed that the −COOH groups on SA, Pec, and TA of the backbone are responsive to pH changes. So the drug loading at pH 7.4 was higher than at pH 1.5 and 5.2. At pH 1.5 the −COOH groups are protonated which contracted the polymer chains by formation of hydrogen bonds and obstruct the release of development of hydrogen bonds and deter the arrival of drug molecules into the networks. At pH 7.4 the −COOH- groups deprotonated to create −COO- anions. The aversions of the adversely charged among the −COO- anions results in an extension of polymer chains henceforth effortlessly allowed to more drug to enter into the networks.

**Fig. 6.**
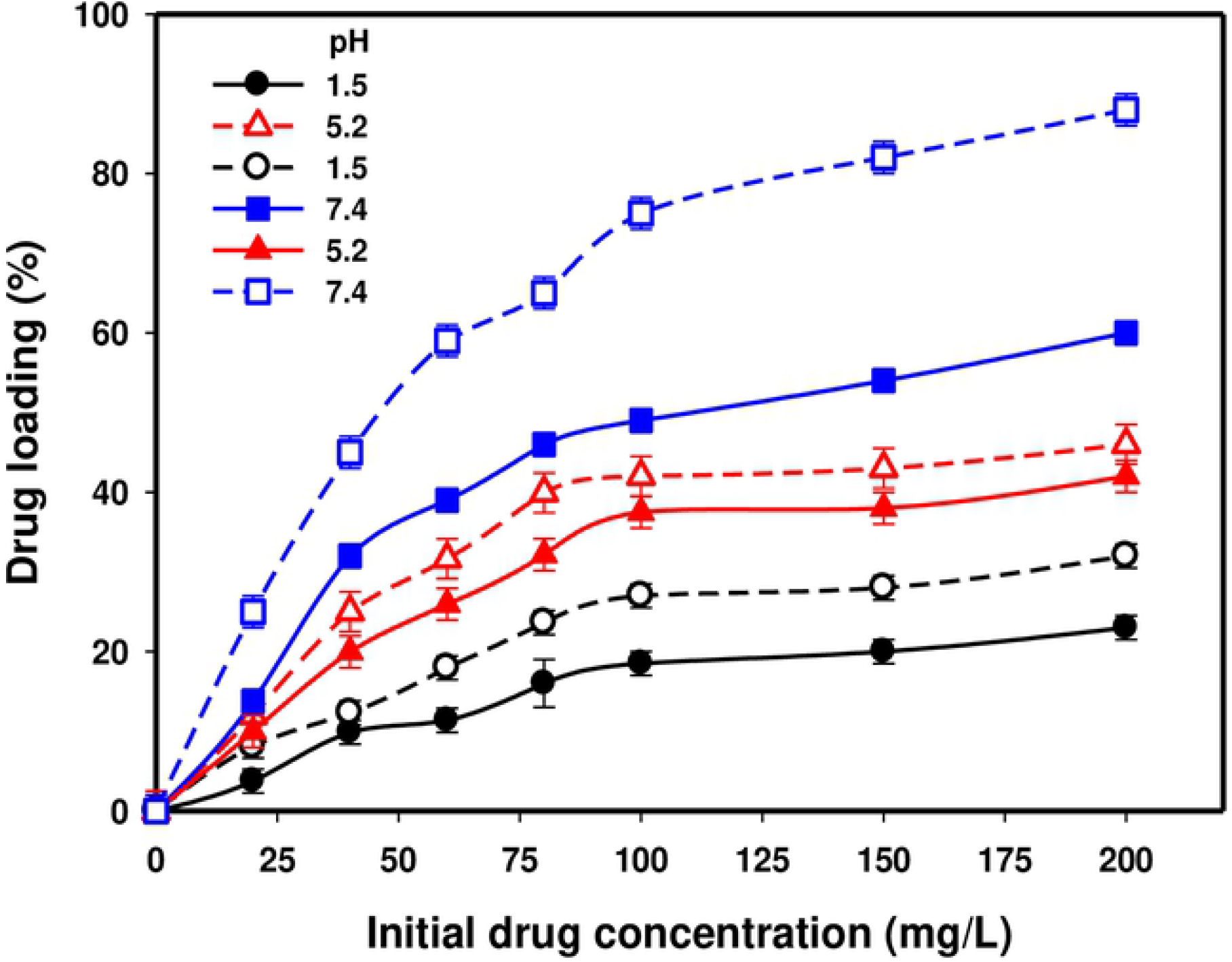
Drug loading of Alg/Pec/TA hydrogel (dark symbols) and Alg/Pec/TA–Ag (open symbols) nanocomposite at different pHs.

### In vitro Propranolol drug release studies

The cumulative percentage of Propranolol release from SA/Pec/TA hydrogel, and SA/Pec/TA-Ag nanocomposite as a function of time at 37°C is shown in Fig. 7. It is clear that the drug release at pH 7.4 was much higher than that at pH 2.1. The drug release from SA/Pec/TA-Ag nanocomposite was 46% at pH 2.1 and was 96% at pH 7.4 within 420 min. Drug releasing behavior relies upon the properties and behavior of matrix stacked this drug. The release of water dissolvable drug from a system happens simply after permeation of water into the matrix which swells and dissolves the drug, trailed by diffusion of the drug [26]. In other meaning, the external medium penetrated into the matrix by the osmotic pressure as well as the drug dissolved and released into the medium. Based on the consideration both of SA/Pec/TA hydrogel, and SA/Pec/TA-Ag nanocomposite dissociated in basic medium (7.4) and associated in acidic medium (2.1) as discussed in the swelling behavior. So the pH sensitive hydrogels are effective drug transporters for a specific site where drug release can be controlled by pH changing. On the other hand, it can be observed that the drug release of SA/Pec/TA-Ag nanocomposite was higher than SA/Pec/TA hydrogel. This means the chemical formulation is highly affected the drug release. The presence of Ag nanoparticles in the network structure of SA/Pec/TA-Ag nanocomposite enhanced the hydrophilicity as well as the drug release. For all of these, the investigated systems particularly SA/Pec/TA-Ag nanocomposite is candidate for the oral drug carrier. It could keep the drug from destroying by acidic gastric fluid.

**Fig. 7.**
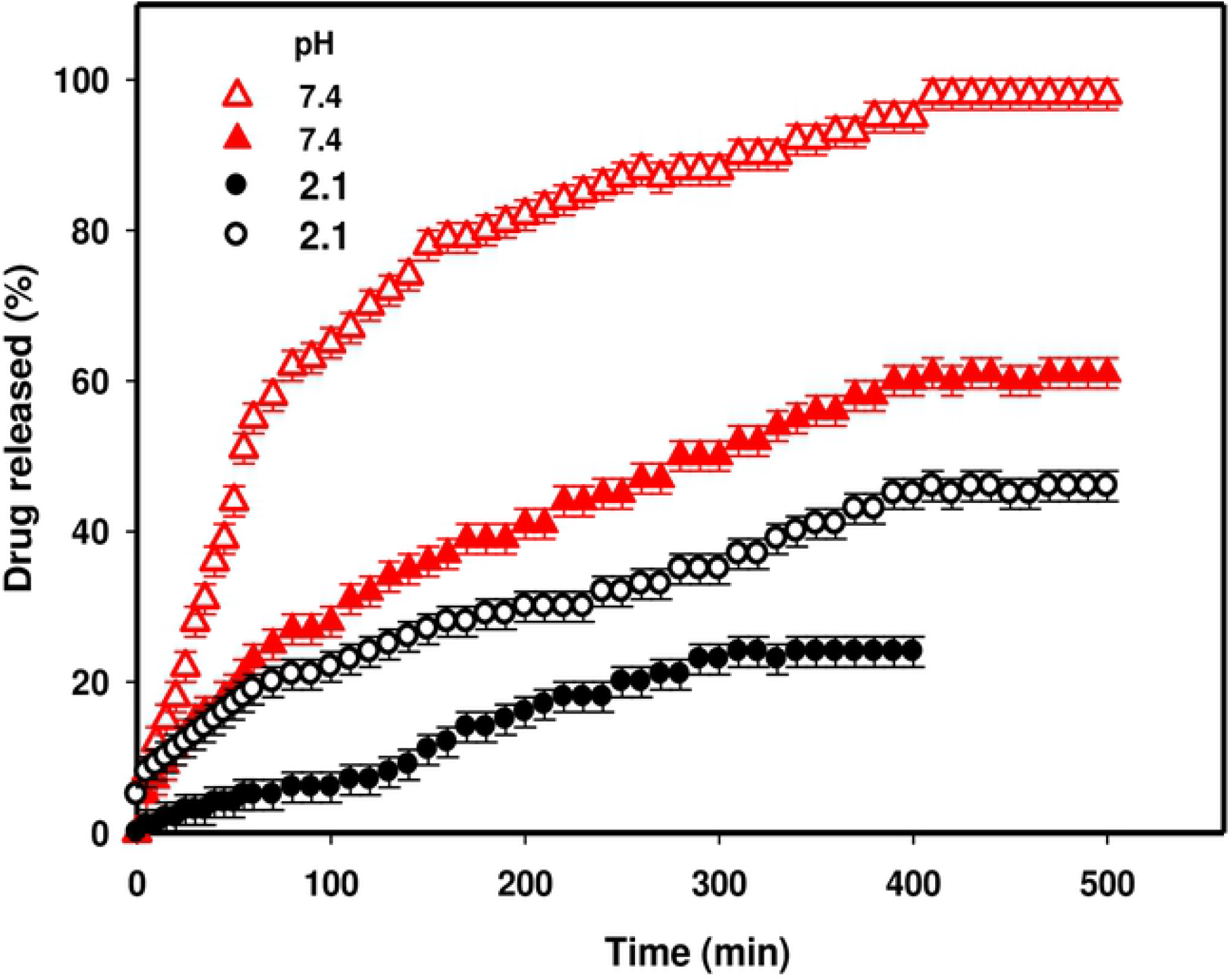
Drug release of Alg/Pec/TA hydrogel (dark symbols) and Alg/Pec/TA–Ag nanocomposite (open symbols) at 37°C.

### Kinetics of releasing

The learning of release kinetics is fundamental for the efficient utilize of the drug carriers. With the end goal to contemplate drug release kinetic mechanism, the information obtained by in vitro release experimental data at pH 7.4 was fitted with different observational kinetic models. These models included zero order, first order, Higuchi Square root, and Ritger-Peppas models. They came about information is appeared in Fig. 8 and the examined information is abridged in Table 1. By contrasting the correlation coefficients, the R^2^ of the Higuchi model is higher than the zero - order and first-order models. This implies the kinetic of Propranolol releasing from SA/Pec/TA hydrogel and SA/Pec/TA-Ag nanocomposite pursues the Higuchi Square root model.

**Fig. 8.**
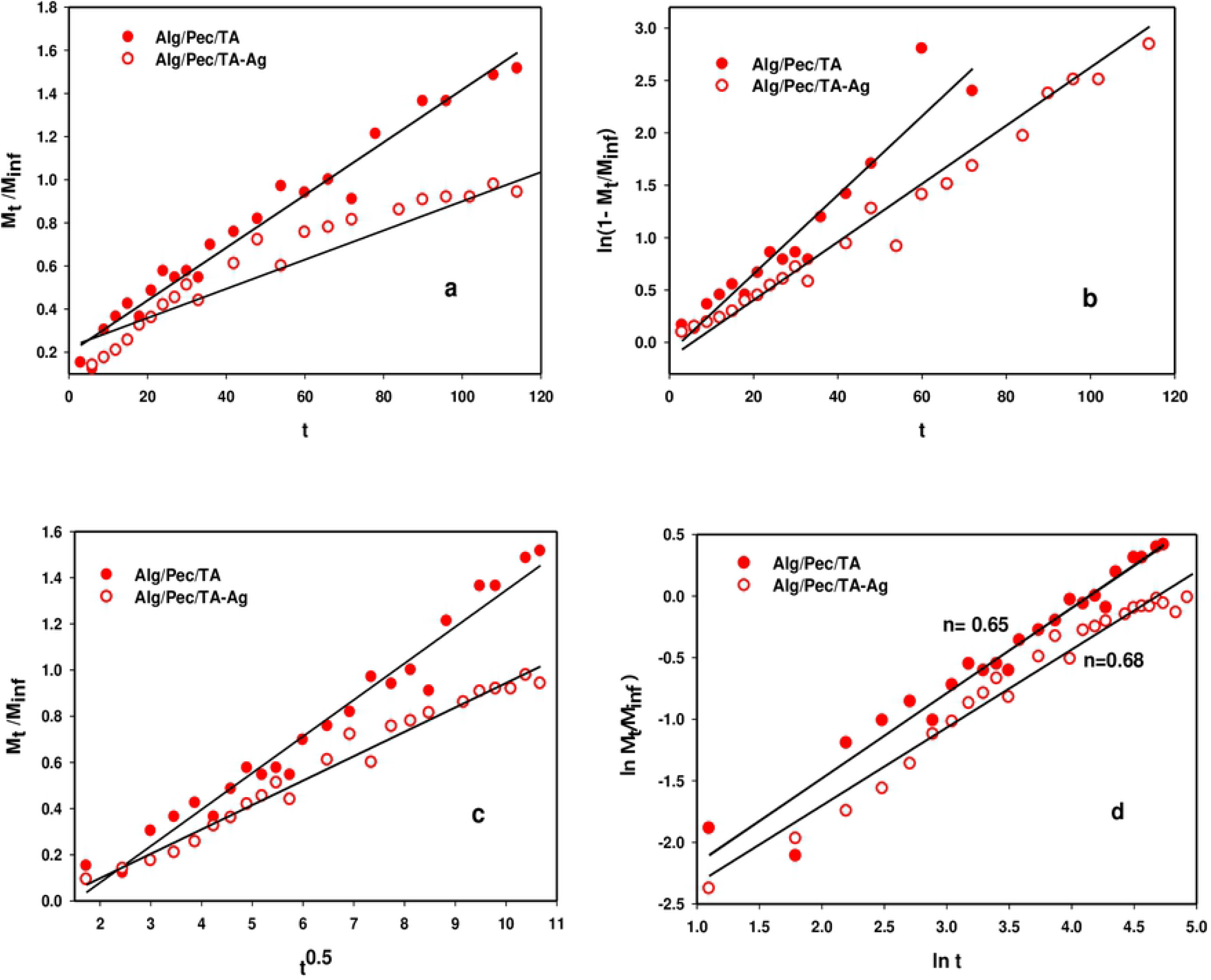
Propranolol release from SA/Pec/TA hydrogel and SA/Pec/TA-Ag nanocomposite fitting curves of different models (a) zero order, (b) first order, (c) Higuchi Square root, and (d) Ritger-Peppas models at 37°C and pH 7.4.

**Table 1.** Kinetic model parameters of different models for Propranolol release from SA/Pec/TA hydrogel and SA/Pec/TA-Ag nanocomposite at 37°C and pH 7.4.

| Variables | SA/Pec/TA | SA/Pec/TA-Ag |
| --- | --- | --- |
| **Zero-order drug release kinetic model** |  |  |
| $R^2$ | 0.9242 | 0.8864 |
| $k_0$ | 0.0122 | 0.0060 |
| **First-order drug release kinetic model** |  |  |
| $R^2$ | 0.9394 | 0.9189 |
| $k_1$ | 0.0278 | 0.0377 |
| **The Higuchi Square root model** |  |  |
| $R^2$ | 0.9679 | 0.9795 |
| $k_1$ | 0.1583 | 0.1057 |
| **The Ritger - Peppas model** |  |  |
| $R^2$ | 0.9542 | 0.9550 |
| k | 2.117 | 5.200 |
| n | 0.6512 | 0.6871 |

For Ritger-Peppas model, the estimation of n is utilized to describe the release mechanism. If n in the range of 0.43 – 0.85 demonstrates non-Fickian release where both diffusion and relaxation control the release. If n ≤ 0.43 shows Fickian release where the release is diffusion control. If n ≥ 0.85 demonstrates case-II transport where the release is relaxation control. Plainly n values for SA/Pec/TA hydrogel and SA/Pec/TA-Ag nanocomposite were in the range of 0.43–0.85. In this way, it tends to be said that the Propranolol release is non-Fickian anomalous transport in which it controlled by both diffusion and relaxation of polymer chains. The release rate of Propranolol from SA/Pec/TA-Ag nanocomposite was higher than SA/Pec/TA hydrogel because of expanding the porosity of the lattice as appeared by surface morphology examination and additionally the swelling capacity thus encouraging the diffusion of the drug.

## Conclusions

A nanocomposite based on Sodium alginate /Pectin/ tannic acid incorporated with silver nanoparticles is synthesized by green method using microwave irradiation. It was found that, the presence of Ag in the nanocomposite confirmed with XRD. The surface morphology totally changed from smooth to a gruff surface by joining of Ag nanoparticles inside the polymeric structure. HRTEM examination explained that an arbitrary dispersion of Ag nanoparticles which showed up as an about circular dark of various particle size average between 21.91 - 34.04 nm by DLS. The swelling studies attained that a higher swelling response of the hydrogel and the nanocomposite attributed to the high hydrophilicity of polymeric chains. pH-sensitivity was confirmed due to the presence of −COOH group and limited swelling obtained at pH 1.5 and higher swelling was done at pH 7.4. The drug release curve demonstrated that the presence of Ag nanoparticles in the network structure of the nanocomposite enhanced the drug release that was 96% at pH 7.4 within 420 min. The drug release was fitted well by Higuchi model and the mechanism found to be non-Fickian anomalous transport. It can be concluded that SA/Pec/TA-Ag nanocomposite is candidate for the oral drug carrier specific for intestinal system.

## Acknowledgments

The authors gratefully acknowledge the Deanship of Scientific Research, Jazan University for the financial support of this work with the project no. JUP8//000322 and also the continuous technical supporting until achieving this work.

